# A Deep Learning Pipeline for Morphological and Viability Assessment of 3D Cancer Cell Spheroids

**DOI:** 10.1101/2025.01.20.633939

**Authors:** Ajay K. Mali, Sivasubramanian Murugappan, Jayashree Rajesh Prasad, Syed A. M. Tofail, Nanasaheb D. Thorat

## Abstract

Three-dimensional (3D) spheroid models have advanced cancer research by better mimicking the tumour microenvironment compared to traditional 2D dimensional (2D) cell cultures. However, challenges persist in high-throughput analysis of morphological characteristics and cell viability, as traditional methods like manual fluorescence analysis are labour-intensive and inconsistent. Existing AI-based approaches often address segmentation or classification in isolation, lacking an integrated workflow.

We propose a scalable, two-stage deep learning pipeline to address these gaps: (1) a U-Net model for precise detection and segmentation of 3D spheroids from microscopic images, achieving 95% prediction accuracy, and (2) a CNN Regression Hybrid method for estimating live/dead cell percentages and classifying spheroids, with an *R*^2^value of 98%. This end-to-end pipeline automates cell viability quantification and generates key morphological parameters for spheroid growth kinetics.

By integrating segmentation and analysis, our method addresses environmental variability and morphological characterization challenges, offering a robust tool for drug discovery, toxicity screening, and clinical research. This approach significantly improves efficiency and scalability of 3D spheroid evaluations, paving the way for advancements in cancer therapeutics.

## 1. Introduction

Three-dimensional (3D) spheroid models have revolutionized cancer research by better mimicking the tumour microenvironment compared to traditional two-dimensional (2D) cultures.^1,2^ These models provide crucial insights into tumour growth kinetics, drug responses, and cellular interactions. However, the complex structure of 3D spheroids and the challenges of fluorescence imaging present significant barriers to automated, high-throughput analysis.^3,4^

Deep learning has emerged as a powerful tool for biological image analysis, offering exceptional performance in segmentation, classification, and quantitative evaluation tasks.^4,5^ Among these, U-Net, a deep learning architecture tailored for biomedical image segmentation, has proven highly effective in extracting regions of interest (ROIs) such as spheroids and nuclei from fluorescence images.^6,7^ U-Net was developed by Ranneberger ^8^ as an effective tool for the segmentation of cellular structures. Its ability to capture fine details and contextual features makes it particularly suitable for 3D spheroid segmentation in fluorescence microscopy.^7,9^ Complementing U-Net, convolutional neural networks (CNNs) have demonstrated robust predictive capabilities, especially when coupled with regression layers for continuous variable estimation, such as live/dead cell percentages in live/dead assays.^10,11,12,13^

Despite these advances, current approaches often address segmentation or viability analysis in isolation, limiting their utility in high-throughput workflows. Traditional threshold-based fluorescence quantification methods remain prone to variability from imaging conditions and user bias.^14,15^ To address these gaps, we propose a two-stage deep learning pipeline that integrates U-Net for precise spheroid segmentation and a CNN Regression model for cell viability prediction. This approach facilitates comprehensive fluorescence image analysis, automates morphological characterization, and ensures scalability for applications like drug discovery and toxicity screening.

Our pipeline addresses critical challenges in spheroid analysis, including variability in growth dynamics, imaging inconsistencies, and the need for reproducible high-throughput evaluations. By integrating segmentation and viability assessment into a unified framework, the proposed method offers a robust, scalable, and automated solution for advancing cancer research.^7,16^

## 2. Materials and Methods

### 2.1 Spheroid Preparation and Live/Dead Assay

3D spheroids were prepared using glioblastoma (U87) and neuroblastoma (SH-SY5Y) cell lines to model tumour heterogeneity. A co-culture method, adapted from prior research, mimicked the tumour microenvironment. Live/Dead assays were performed using Calcein-AM (green fluorescence for live cells) and Ethidium Homodimer-1 (red fluorescence for dead cells). Samples were incubated for 30 minutes at 37°C before excess dye was removed.^17^ This combination provides a robust in vitro model to replicate glioblastoma’s complex cellular interactions.

The Live/Dead Assay was used to evaluate the spheroid’s cell viability. Viable cells were separated from those with affected cells using Calcein-AM, which emits green fluorescence to identify live cells, and Ethidium Homodimer-1, which emits red fluorescence to show dead cells. The samples were incubated with the dyes for 30 minutes at 37°C as part of the staining process, and any excess dye was then washed off. The used method ensures enhanced precision in assessing the viability of cells within the spheroid. ^17^

### 2.2 Image Acquisition and Preprocessing

Fluorescence images were captured using a confocal microscope and resized to 128×128 pixels to reduce computational complexity while preserving morphological features. Standardization of image dimensions minimized variability and facilitated consistent training for the deep learning models.^18,19^ By lowering image size, researchers may regulate the additional storage space and processing time necessary for larger images, allowing for more effective analysis in limited in resources situations.^20^ In addition, standardization helps in limiting the variability produced by various image sizes, hence preserving significant characteristics required for accurate feature extraction.

### 2.3 Spheroid Segmentation Using U-Net

The U-Net model was employed for spheroid segmentation. Training used 500 annotated fluorescence images, with segmentation masks created in ImageJ. Key hyperparameters included a learning rate of 0.001, 20 epochs, and binary cross-entropy loss. Data augmentation (rotation, flipping, intensity scaling) enhanced generalization.^21,22,23^ post-segmentation, morphological filtering was applied to isolate regions of interest (ROIs).^13^

The workflow for the U-Net model as shown in Fig.1

**Fig 1.**
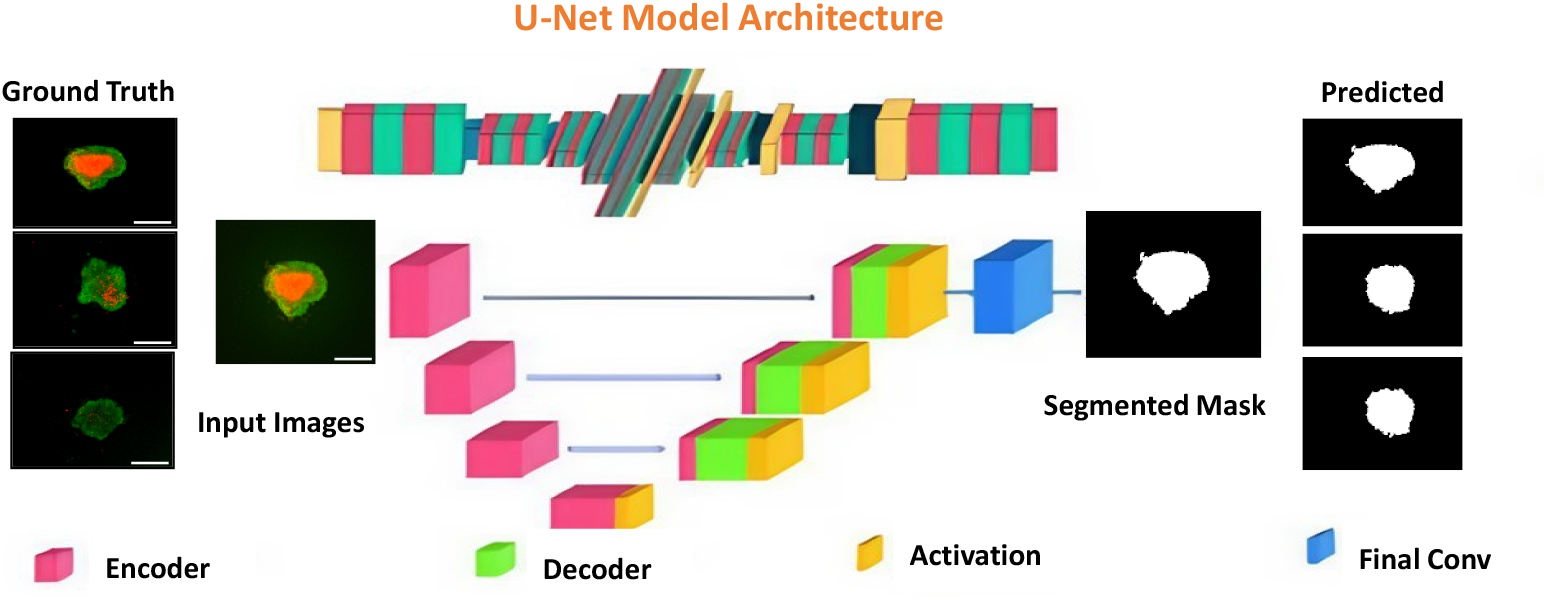
The above flowchart demonstrates the U-Net model procedure. It starts with an input image, then the encoder path (down sampling), bottleneck, and decoder path (up sampling), which includes skip connections to retain spatial information for accurate segmentation.

### 2.4 Cross-Validation Technique

To improve model performance and prevent overfitting, K-fold cross-validation (K=10) was implemented. This involved dividing the dataset into ten separate folds, iteratively training on nine folds and validating on the remaining fold, so providing an accurate evaluation of model generalization across various data subsets.^24^ The accuracy of predictions was assessed using measures like the Coefficient of Determination, R^2^, and Mean Absolute Error (MAE). The model demonstrated significant predictive performance by achieving a high R^2^. value of 97% on the test datasets. The model’s performance consistency and predictability are further demonstrated by the MAE graph across folds and the accuracy and loss graphs for each fold (see Supplementary Figs. 1, 2, 3).

### 2.5 Viability Prediction Using CNN + Regression

The segmented ROIs were input to a CNN-based regression model to predict live and dead cell percentages. The CNN architecture included convolutional layers with ReLU activation, max-pooling, and a regression output layer. Ground truth viability data were derived from fluorescence intensity ratios of Calcein-AM and Ethidium Homodimer-1 channels. A 10-fold cross-validation method was used to split the dataset into training and testing subsets, providing an extraordinary predictive accuracy with a mean R^2^ value of across all fold are 89%. The workflow for the CNN Regression hybrid model demonstrated in Fig.2

**Fig 2.**
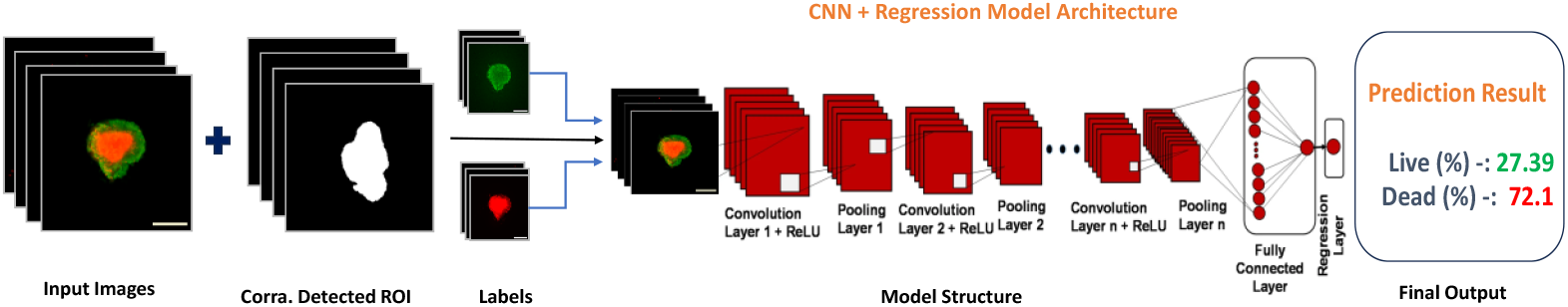
The CNN + Regression model procedure is illustrated by the flowchart above. The CNN collects relevant spatial information from the input image and passes them to a regression layer, which predicts continuous output of live and dead cell percentages.

### 2.6 Morphological Analysis

Morphological metrics, such as spheroid area, sphericity, and roundness, were measured using Python libraries (scikit-image, OpenCV) and ImageJ. ^25^ Results were cross-validated, showing Python-based methods to be more efficient and consistent.^26,27^

### 2.7 Evaluation Matrices

For segmentation, Dice Similarity Coefficient (DSC) and Intersection Over Union (IoU) quantified overlap between predicted and ground truth masks. For viability prediction, R^2^ and MAE assessed model performance. These metrics ensured balanced evaluation across imbalanced datasets and provided insights into prediction accuracy.

#### For Spheroid Segmentation (U-Net Model)

1. **Dice Similarity coefficient -** The U-Net model’s performance was assessed using the Dice Similarity Coefficient (DSC) to quantify the overlap between the predicted segmentation (P) and the ground truth (G) masks. The DSC is defined as:

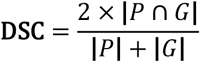 The Dice coefficient is extensively utilized in medical image analysis due to its ability to demonstrate consistency in overlapping regions, making it particularly successful for segmentation tasks. This is particularly crucial for situations such as spheroid segmentation, when precision in border identification is critical. Unlike pixel accuracy, DSC penalizes both false positive and false negatives, ensuring a balanced evaluation for highly im-balanced datasets typical in medical imaging.^28,29^
2. **Intersection Over Union (IoU**) - Another overlap-based metric, measures the ratio of the intersection to the union of the predicted and ground truth masks:

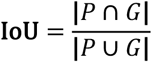 IoU enhances the DSC by providing a More comprehensive assessment, particularly sensitive to the presence of false positives. It is often used in medical image analysis because it can constantly evaluate both small and large frameworks, making sure that segmentation works well across a range of spheroidal sizes and imaging conditions.^28,29^

### For Live/Dead Viability Prediction (CNN + Regression Model)

1) **Coefficient of Determination (*R*^2^)-** The *R*^2^metric, or the coefficient of determination, assess how well the predicted values 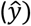 approximately the actual value (*y*). It is mathematically expressed as:

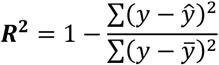
  - 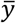: Mean of actual values.
  - An *R*^2^ close to 1 reflects excellent predictive performance captures the ratio of variability explained by the model, giving insights into its predicative power.^30^
3) **Mean Absolute Error (MAE) -** Measures the average absolute differences between predicted and actual values.

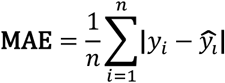 Where:
  - *y*_*i*_= actual values
  - 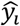 = predicted values
  - *n*= number of data points

MAE is widely used in regression tasks to assess model performance by quantifying the magnitude of errors. For live/ dead prediction, MAE provides a simple interpretation of average prediction errors, allowing researchers to identify practical limitations in viability quantification.^31^

### 2.8 Computational Setup

Models were developed using Python libraries (TensorFlow, Keras, scikit-image) and trained on an NVIDIA RTX 3090 GPU. The computational pipeline was optimized for resource efficiency without compromising accuracy.

## 3. Results

### 3.1 U-Net Spheroid Detection and Segmentation Accuracy

In this segment, we demonstrate the efficacy of the U-Net model for spheroid detection and segmentation. To evaluate precision, the model’s segmentation outputs were juxtaposed with manually annotated ground truth masks. The U-Net model excels at segmenting spheroids characterized by clear boundaries and uniform fluorescence intensity, achieving high accuracy even in images with minimal background noise or distinctly defined spheroids. However, complexity occur in images with overlapping spheroids or significant background objects, leading the model to sometimes misclassify areas or insufficiently define boundaries. These occurrences underscored the necessity for sophisticated preprocessing methods or supplementary training data to enhance segmentation precision in these complex scenarios. Despite those challenges, the model’s general robustness was apparent from its constant Dice coefficient of 95% and precision of 98% (Supplementary Fig.4) in detecting spheroid regions from the background. Segmentation accuracy was evaluated using metrics including the Dice coefficient, Intersection over Union (IoU), and pixel-wise accuracy (Supplementary Fig. 4), demonstrating the model’s proficiency in reliably segmenting spheroids across diverse image datasets. The segmentation masks generated by the U-Net model closely matched the ground truth, confirming the model’s robustness and reliability.

**Fig 3.**
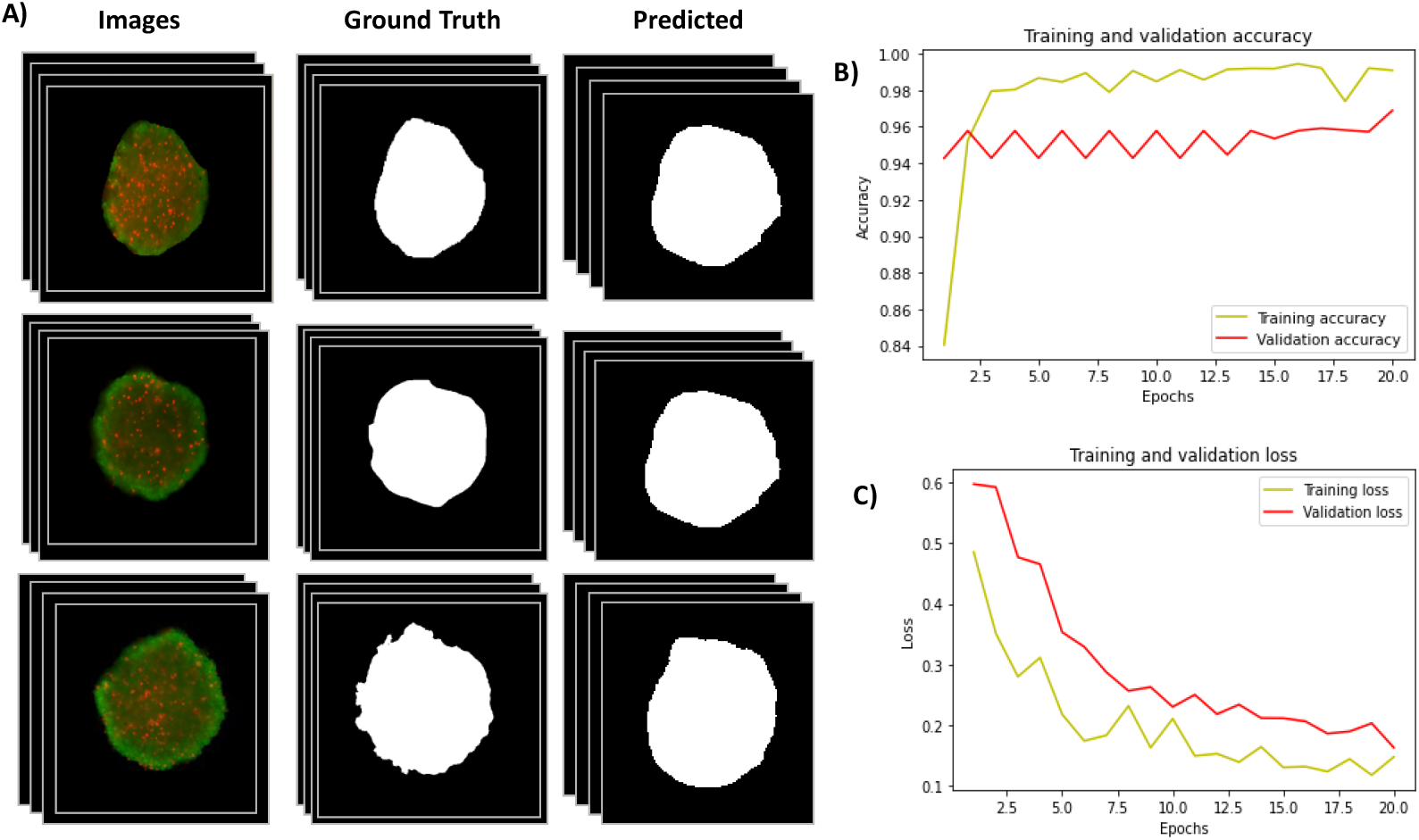
Results of spheroid segmentation using the U-Net model. A) The left column shows the original fluorescence images, followed by the corresponding ground truth masks (manually annotated using ImageJ) and the predicted segmentation masks by the model. The quantitative assessment revealed a Dice coefficient of 95%, signifying considerable overlap between the expected and actual masks. The training and validation performance curves demonstrate the model’s efficacy in learning and generalizing spheroid segmentation. The training accuracy steadily improved to a peak of 98%, while validation accuracy consistently above 96%, demonstrating the model’s high precision across datasets. The loss curves demonstrate consistent advancement, with training loss decreasing to roughly 0.05 and validation loss stabilizing at around 0.1 after 20 epochs. This indicates negligible overfitting and validates the model’s strong ability to segment spheroids with high reliability. The persistent disparity between training and validation loss indicates the model’s capacity to generalize proficiently to unfamiliar data.

**Fig 4.**
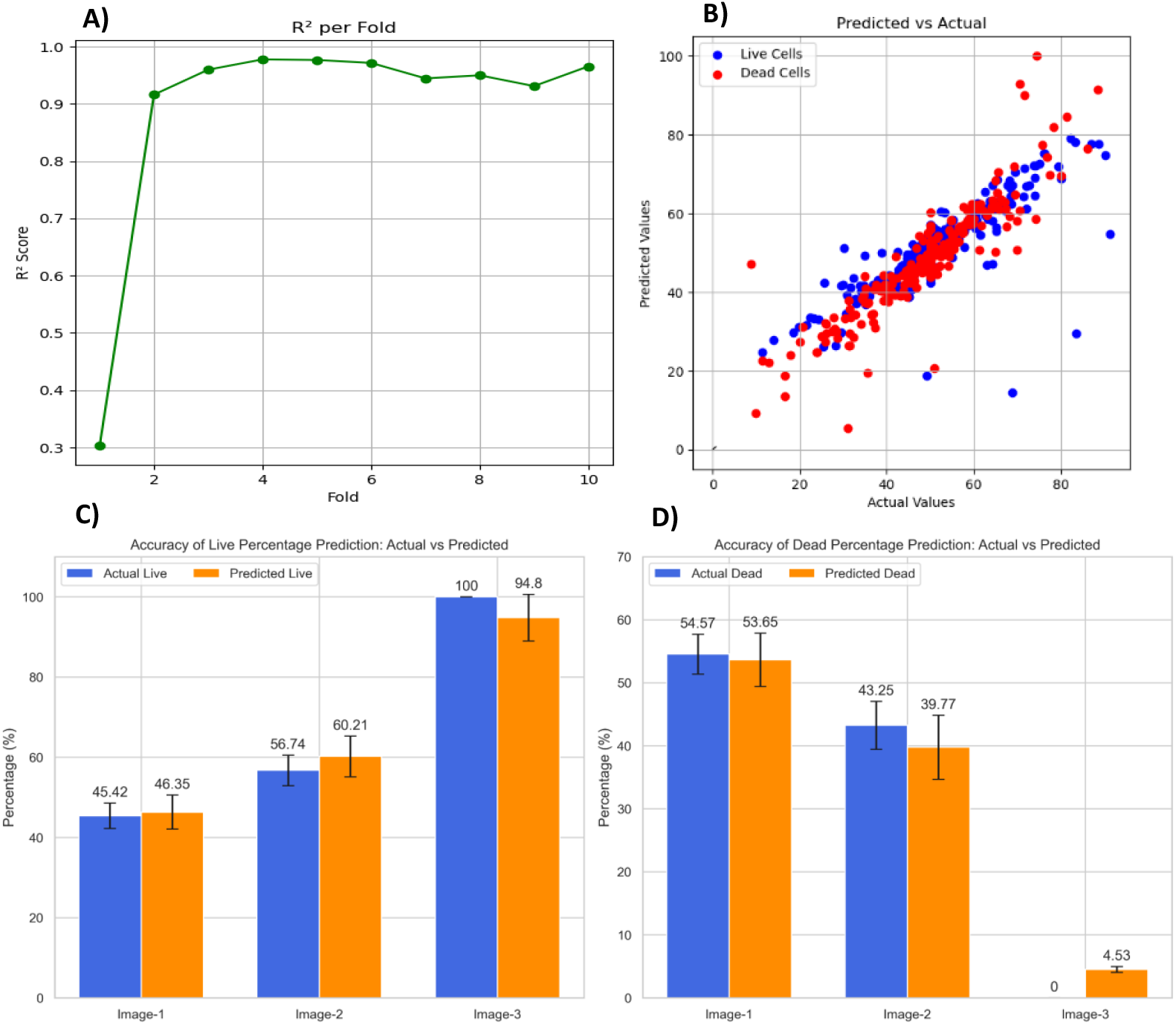
A) R^2^ per Fold Across 10-Fold Cross-Validation: The R2 values obtained during cross-validation are displayed in the plot. Except for a marginally lower result in Fold-1, the model consistently obtains excellent R2 scores (0.91–0.98) across all folds. This shows how well the model predicts viability percentages with the least amount of error, demonstrating its generalizability and robustness. B) **The scatter plot of actual vs. predicted percentages** shows a close alignment of data points along the diagonal line, indicating a significant positive correlation between the two. This demonstrates how well the model predicts the percentages of live and dead cells under various experimental settings. C) **Accuracy of Live Percentage Prediction** and D) **Dead Percentage Prediction:** The bar charts illustrate the actual and predicted percentages of viable and non-viable cells for three spheroid images, with error bars denoting the associated errors. The CNN-based regression model has significant accuracy in aligning predictions with manual counts. Image-3 distinctly demonstrates the model’s capacity to accurately forecast extreme scenarios (e.g., 100% live and 4.53% dead).

### 3.2 Viability Prediction Outcomes

The CNN-based regression model quantified the proportions of live and dead cells in the segmented spheroid, with predictions validated by manual cell counts derived from fluorescence intensity ratios of Calcein-AM (live cells) and Ethidium Homodimer-1 (dead cells). The results, illustrated in the “Accuracy of Live Percentage Prediction” and “Accuracy of Dead Percentage Prediction” graphs, exhibit a strong correlation between the actual and predicted values for both live and dead cell percentages. In Image-3, the predicted live proportion (94.8%) closely matches the actual live percentage (100%), while the expected dead percentage (4.53%) aligns with the very negligible actual dead percentage, highlighting the model’s precision in extreme scenarios.

Across other images, the model performs consistently with low mistakes. Notably, the error bars in the graphs indicate confidence intervals, which helps to validate the forecasts’ robustness. The R^2^ score of 0.98 indicates a great fit between predicted and actual values, capturing 98% of the variability in the data.

The second graph, “R^2^ per Fold,” illustrates the model’s stability and generalizability during cross-validation. The R^2^ values consistently remain elevated across 10 folds, varying from 0.91 to 0.98. The initial fold (Fold-1) is markedly lower at 0.31. The variance in the initial fold is possibly due to data partitioning, potentially implemented in challenging scenarios during that fold. The overall trend of R^2^ values indicates a well-trained and dependable model that can generalize new data.

The model has good predictive accuracy, however there are slight gaps, such as a ±4% error in live percentage forecasts in Image-2 and a lower R^2^ value of 0.31 in Fold-1 during cross-validation, which may be attributed to dataset splitting. These constraints indicate areas for improvement, such as increasing the dataset or improving resilience with additional regularization techniques.

These findings highlight the model’s capability to automate live-dead cell classification tasks with high precision, notably reducing the necessity for manual involvement. This automation can enhance throughput in high-content drug screening and enable accurate viability evaluation under different experimental conditions, opening the path for its implementation in both preclinical and clinical research settings.

### 3.3 Morphological Analysis Findings

The morphological analysis concentrated on the spheroid’s growth kinetics, how culturing density affected spheroid development, and how morphological metrics changed over time. Both automatic Python-based techniques and manual ImageJ analysis were used to quantify spheroid area, sphericity, and roundness. The findings showed that the spheroid grew larger over time, and that the initial culturing density had a significant impact on the growth rate. Increased spheroid volume and quicker aggregation were the results of higher seeding densities (Fig.5). While the ImageJ-based method had somewhat higher variability because of manual intervention, the Python-based approach using OpenCV and scikit-image produced more consistent and efficient measurements, according to a comparison of manual and AI-based methods.

The plots highlight the impact of initial culturing density on spheroid size by comparing the growth of MCF-7 spheroid over a period of 14 days. Three different seeding densities (4000, 6000, and 8000 cells) were evaluated, and their growth kinetics were monitored by measuring spheroid area.

**Fig 5.**
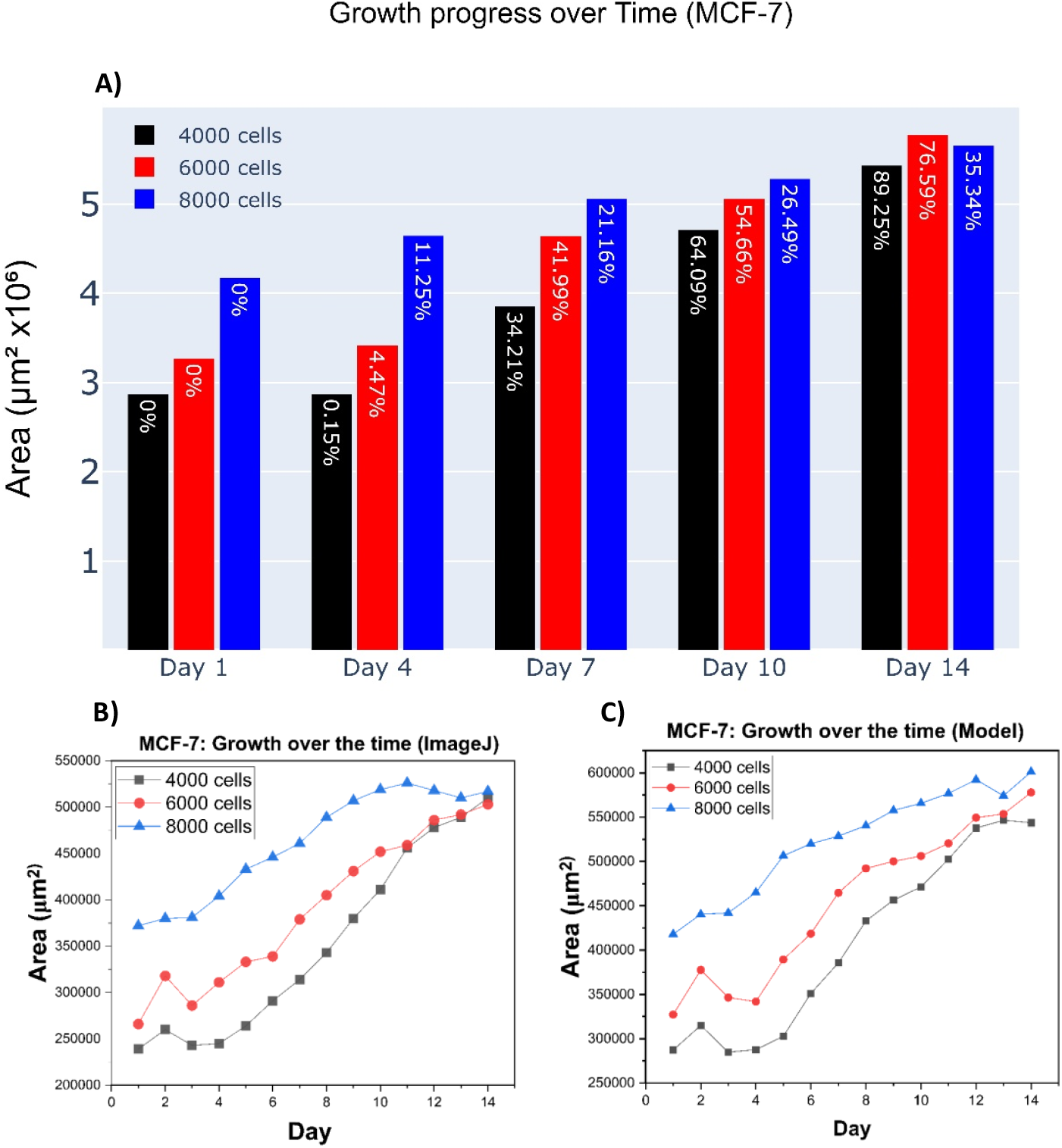
A) Growth Kinetics of MCF-7 Spheroid. The graphs show the spheroid growth in terms of area (in µm^2^ × 10^6^) over 14 days for three different seeding densities (4000, 6000, and 8000 cells B) This graph shows the area of the spheroid over 14 days, calculated using manual tool ImageJ. C) The graph shows the spheroid’s area over a period of 14 days, as determined by our model technique. Significant parallels between the two graphs become noticeable when we look more closely.

### 3.4 Application in Drug Development and Toxicology

To demonstrate the AI-based system’s potential in drug development and toxicity testing, we ran case studies with simulated drugs. Spheroids were treated to TMZ at different concentrations (10 µM, 100 µM, and 1000 µM) and viability changes were observed. The AI system enables high-throughput screening by immediately predicting cell viability and quantifying drug treatment responses. The model was also useful in identifying dangerous chemicals, as it recognized significant cell decreases at lower drug doses, providing information on potential cytotoxic effects. These findings demonstrated how the AI system may speed up the drug discovery process, increasing efficiency and accuracy in detecting prospective drug candidates and toxicological potential dangers. When we compare this Artificial intelligence-enhanced results with Manual results it is showing notably similar results to that.

The live/dead assay visualized in Fig. 6 shows the impact of different concentrations of a TMZ drug on cellular viability in two different cell lines (U87MG and SH-SY5Y) as well as their co-culture. Images indicate that at lower drug concentrations (10 μM and 100 μM), live cells are still common throughout treatment groups, demonstrating good cell viability. Green fluorescence indicates a healthy number of live cells in the control group, while red-stained dead cells are minimal. However, at the greatest dose (1000 μM), red fluorescence increases significantly, especially in the SH-SY5Y line, indicating increased cell death.

**Fig 6.**
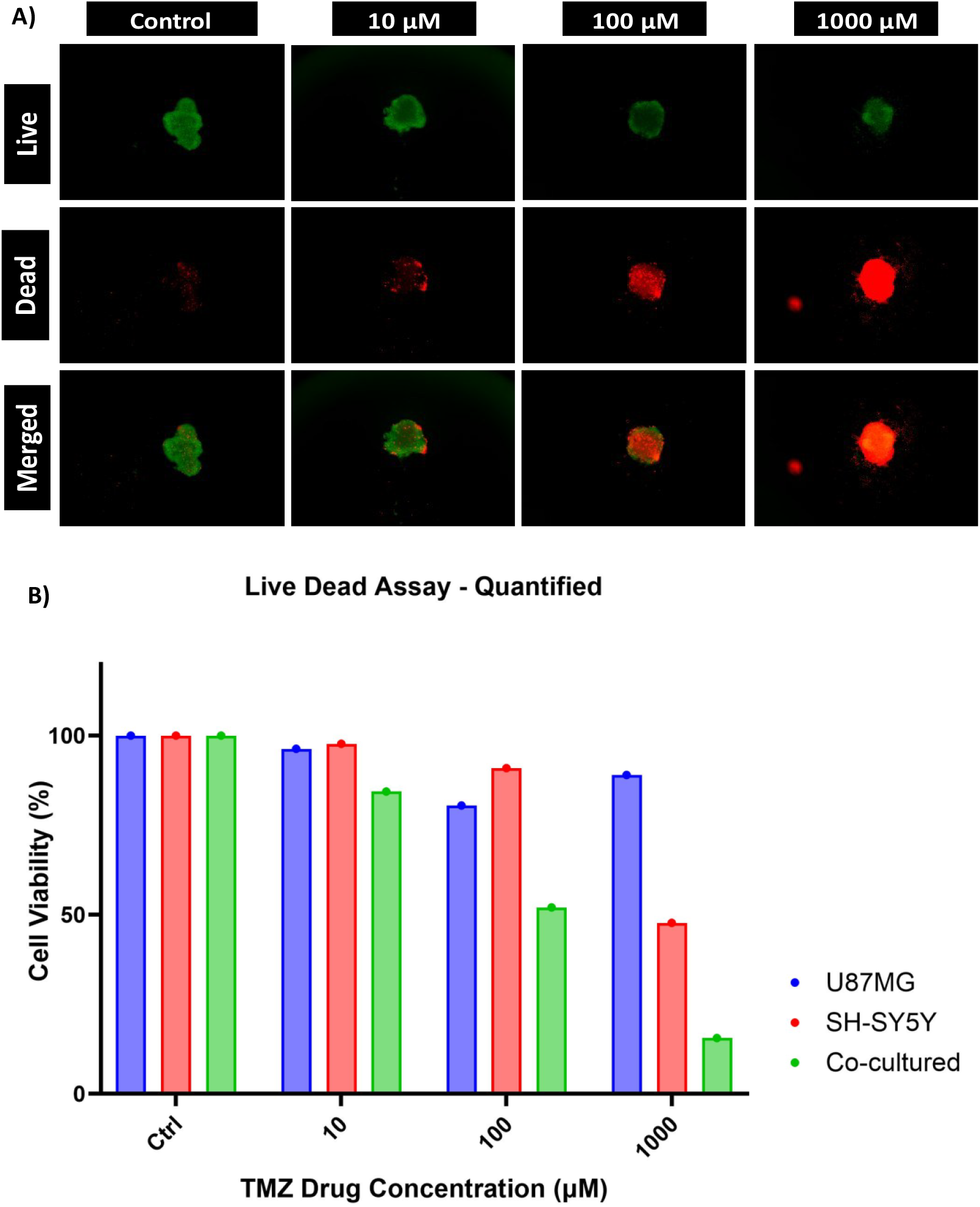
A) Live/Dead Assay Results. Images from fluorescent microscopy that display both living (green) and dead (red) cells at various TMZ concentrations (Control, 10 μM, 100 μM, and 1000 μM). Live cells are shown in the top row, dead cells are shown in the middle row, and the combined findings are displayed in the bottom row. Dead cell fluorescence increases with increasing TMZ doses, especially at 1000 μM. B) **Live Dead Assay - Quantified**. Cell viability (in percentage) of co-cultured cells (green), U87MG (blue), and SH-SY5Y (red) in response to increasing TMZ concentrations (0 (Ctrl), 10, 100, and 1000 μM) is shown in a bar graph. According to statistical analysis, all cell lines exhibit high viability at lower concentrations (0 (Ctrl), 10 μM, and 100 μM; however, SH-SY5Y cells show a notable decrease at 1000 μM.

Figure 6 presents quantitative analysis of live/dead assay visuals. Results show that cell viability in U87MG and co-cultured cells remain almost 100% at doses of 10 μM and 100 μM. Viability considerably decreases at 1000 μM, especially in the SH-SY5Y cell line, compared to other groups. The dose-dependent cytotoxic effect of TMZ shows that different cell lines respond differently to treatment.

## 4. Discussion

The application of AI-based systems for 3D spheroid analysis line up with standards set in the field of biomedical research. Research has demonstrated that high-throughput screening enhanced by artificial intelligence (AI) can markedly decrease the duration needed to identify lead compounds for drug discovery. Three-dimensional spheroid models replicate in vivo-like circumstances and can now be analysed more efficiently and effectively using AI algorithms.^32,33^

As noted by developments in regenerative medicine have also employed computerized analysis to measure morphological changes in response to treatments.^34^ This speeds up the creation of tissue-engineered models and increases reproducibility.

In personalized medicine, studies such as by^35,36^ Emphasize the utility of patient-derived spheroids in developing personalized medication sensitivity profiles. The analysis conducted using artificial intelligence improves this process by delivering accurate and quick evaluations, which are essential in reducing rely on animal models while maintaining standards of ethics.

In comparison to conventional human techniques, artificial intelligence spheroid analysis provides substantial enhancements in accuracy, consistency, and time efficiency. Manual techniques, including visual examination and manual counting, are prone to human error and subjectivity, resulting in diversity in outcomes, particularly when analyzing extensive datasets or intricate 3D structures. Conversely, the AI-driven system provides uniform, replicable outcomes, minimizing operator bias and reducing inter-observer variability.

The AI-based model exhibited a significant connection with manual cell viability counts, evidenced by a robust R^2^ value between predicted and actual data. Moreover, the AI system facilitates the identification of small morphological alterations in spheroids that may be overlooked by manual techniques, so offering a more thorough comprehension of cellular responses to treatments. The automated characteristics of the artificial intelligence (AI) technology significantly enhance time efficiency. The model can analysed hundreds of spheroids in a fraction of the time required for a researcher to manually count cells or evaluate morphological features, providing it extremely ideal for high-throughput applications.

Despite it was clear benefits, the artificial intelligence-driven spheroid analysis system possesses limits that want attention in future work. A main problem is to potential complications with dye penetration during the Live/Dead experiment. In thick 3D spheroid, the ability of Calcein-AM and Ethidium Homodimer-1 to penetrate consistently can be limited, leading to inaccuracies in viability assessments, particularly in the core of larger spheroid. Incomplete staining may result in ignoring the proportion of dead cells in the centre of the spheroid. Advancements in dye compositions or the incorporation of multi-staining techniques might reduce this limitation by enhancing the spatial resolution of viability and functional evaluations.

A further limitation occurs from the imaging difficulties caused by confocal microscopy, particularly when capturing images of large or dense spheroids. The quality of fluorescence images can be influenced by issues including light scattering, optical aberrations, and depth penetration, especially when imaging deeper areas of the spheroid. Advancements in imaging methods, such light-sheet microscopy and enhanced fluorescent probes, may enhance image quality and address some problems, facilitating more precise study of deeper spheroid regions.

Future advancements in multi-staining techniques may enhance the model’s capacity to evaluate both cell viability and functionality simultaneously. Multi-channel imaging, utilizing diverse biomarkers to evaluate multiple facets of cellular health (e.g., proliferation, apoptosis, and necrosis), would yield a more thorough comprehension of spheroid behaviour. This would be especially beneficial in drug screening, where it is crucial to understand not only cell viability but also the unique processes of drug-induced toxicity or therapeutic response.

The implementation of AI-driven spheroid analysis in healthcare environments presents ethical and practical challenges. Maintaining patient data confidentiality and reducing biases in model predictions are essential for balanced results. Compliance with rules like as GDPR and the rigorous assessment of artificial intelligence (AI) models for clinical-grade integrity are crucial. Practically, the capacity for high-throughput workflows, the standardization of imaging and staining methods, and sufficient training for physicians are essential measures for ensuring successful integration into clinical practices. These approaches will facilitate the system’s broad acceptance while preserving ethical purity and operational efficiency.

## 5. Conclusion

This study presents an innovative Artificial intelligence-based framework for the analysis of 3D spheroid models, which is significant advancement in cancer research. The two-stage pipeline integrates a U-net model for accurate segmentation of 3D spheroids, achieving a prediction accuracy of 95%, and a CNN Regression Hybrid approach for quantifying live/dead cell percentages, with an R^2^ value of 98%. This integration of segmentation and viability analysis addresses key challenges in high-throughput evaluations, including environmental variability and morphological characterization. By automating this process, the system enhances scalability, efficiency, and precision, cover the way for its application in drug discovery, toxicity screening, and clinical research.

Looking ahead, addressing challenges like dye penetration variability and imaging inconsistencies will enhance the robustness of this pipeline. Adding biomarkers and adopting multimodal AI models can provide deeper insights for complex cell behaviours. Advances in Ai techniques, such as self-supervised learning and generative approaches, offer potential to uncover complicated biological patterns. By encouraging cross-disciplinary collaborations, this framework is assured to direct innovations in personalized medicine, therapeutic development, and translational research.

## Notes

### Competing Interest Statement

The authors have declared no competing interest.

## References

1. Gheytanchi, E. et al. Morphological and molecular characteristics of spheroid formation in HT-29 and Caco-2 colorectal cancer cell lines. Cancer Cell Int 21, 1–16 (2021).

2. Manduca, N., Maccafeo, E., De Maria, R., Sistigu, A. & Musella, M. 3D cancer models: One step closer to in vitro human studies. Front Immunol 14, 1175503 (2023).

3. Chen, Z. et al. Automated evaluation of tumor spheroid behavior in 3D culture using deep learning-based recognition. Biomaterials 272, (2021).

4. Shirai, K. et al. The importance of scoring recognition fitness in spheroid morphological analysis for robust label-free quality evaluation. Regen Ther 14, 205–214 (2020).

5. Zhou, S. K. et al. A Review of Deep Learning in Medical Imaging: Imaging Traits, Technology Trends, Case Studies with Progress Highlights, and Future Promises. Proceedings of the IEEE 109, 820–838 (2021).

6. Weng, W. & Zhu, X. INet: Convolutional Networks for Biomedical Image Segmentation. IEEE Access 9, 16591–16603 (2021).

7. Çiçek, Ö., Abdulkadir, A., Lienkamp, S. S., Brox, T. & Ronneberger, O. 3D U-Net: Learning Dense Volumetric Segmentation from Sparse Annotation. (2016).

8. Weng, W. & Zhu, X. U-Net: Convolutional Networks for Biomedical Image Segmentation. IEEE Access 9, 16591–16603 (2015).

9. Esteva, A. et al. Deep learning-enabled medical computer vision. npj Digital Medicine vol. 4 Preprint at 10.1038/s41746-020-00376-2 (2021).

10. Grexa, I. et al. SpheroidPicker for automated 3D cell culture manipulation using deep learning. Sci Rep 11, (2021).

11. Huang, G., Liu, Z., Van Der Maaten, L. & Weinberger, K. Q. Densely Connected Convolutional Networks. https://github.com/liuzhuang13/DenseNet.

12. Liu, T., Siegel, E. & Shen, D. Deep Learning and Medical Image Analysis for COVID-19 Diagnosis and Prediction. Annu Rev Biomed Eng 24, 179–201 (2022).

13. Liu, T., Siegel, E. & Shen, D. Deep Learning and Medical Image Analysis for COVID-19 Diagnosis and Prediction. Annu. Rev. Biomed. Eng. 2022 24, 179–201 (2024).

14. Leiwe, M. N. et al. Automated neuronal reconstruction with super-multicolour Tetbow labelling and threshold-based clustering of colour hues. Nat Commun 15, (2024).

15. Pereira, A. R. et al. Development, Validation, and Implementation of an Augmented Multiwell, Multitarget Quantitative PCR for the Analysis of Human Papillomavirus Genotyping through Software Automation, Data Science, and Artificial Intelligence. Journal of Molecular Diagnostics 26, 781–791 (2024).

16. Chieregato, M. et al. A hybrid machine learning/deep learning COVID-19 severity predictive model from CT images and clinical data. Sci Rep 12, (2022).

17. Murugappan, S., Mali, A. K., Tofail, S. A. M. & Thorat, N. D. Astrocyte-Neuron co-cultured 3D tumor spheroid model for Anti-cancer Drug Screening. (2024) doi:10.1101/2024.11.11.622957.

18. Shen, D., Wu, G. & Suk, H.-I. Deep Learning in Medical Image Analysis. 25, 24 (2024).

19. Murcia-Gómez, D., Rojas-Valenzuela, I. & Valenzuela, O. Impact of Image Preprocessing Methods and Deep Learning Models for Classifying Histopathological Breast Cancer Images. Applied Sciences (Switzerland) 12, (2022).

20. Wang, S., Yang, D. M., Rong, R., Zhan, X. & Xiao, G. Pathology Image Analysis Using Segmentation Deep Learning Algorithms. American Journal of Pathology vol. 189 1686–1698 Preprint at 10.1016/j.ajpath.2019.05.007 (2019).

21. Yang, S. et al. Image Data Augmentation for Deep Learning: A Survey. (2022).

22. Deep Learning - Ian Goodfellow, Yoshua Bengio, Aaron Courville - Google Books. https://books.google.ie/books?hl=en&lr=&id=omivDQAAQBAJ&oi=fnd&pg=PR5&dq=Goodfellow,+I.,+Bengio,+Y.,+%26+Courville,+A.+(2016).+Deep+Learning.+MIT+Press.&ots=MOP1iokGSR&sig=q9N6OApTHqVAGWh8CnyG0Gd2xWk&redir_esc=y#v=onepage&q=Goodfellow%2C%20I.%2C%20Bengio%2C%20Y.%2C%20%26%20Courville%2C%20A.%20(2016).%20Deep%20Learning.%20MIT%20Press.&f=false.

23. Kingma, D. P. & Ba, J. Adam: A Method for Stochastic Optimization. (2014).

24. James, G., Witten, D., Hastie, T., Tibshirani, R. & Taylor, J. An Introduction to Statistical Learning. (2023) doi:10.1007/978-3-031-38747-0.

25. Rueden, C. T. et al. ImageJ2: ImageJ for the next generation of scientific image data. BMC Bioinformatics 18, (2017).

26. Sirenko, O. et al. High-Content Assays for Characterizing the Viability and Morphology of 3D Cancer Spheroid Cultures. Assay Drug Dev Technol 13, 402–414 (2015).

27. Ferraro, R., Di Franco, J., Caserta, S. & Guido, S. The morphology of cell spheroids in simple shear flow. Front Phys 12, (2024).

28. Taha, A. A. & Hanbury, A. Metrics for evaluating 3D medical image segmentation: Analysis, selection, and tool. BMC Med Imaging 15, (2015).

29. Milletari, F., Navab, N. & Ahmadi, S.-A. V-Net: Fully Convolutional Neural Networks for Volumetric Medical Image Segmentation. (2016).

30. Kuhn, M. & Johnson, K. Applied predictive modeling. Applied Predictive Modeling 1–600 (2013) doi:10.1007/978-1-4614-6849-3/COVER.

31. Willmott, C. J. & Matsuura, K. Advantages of the mean absolute error (MAE) over the root mean square error (RMSE) in assessing average model performance. Clim Res 30, 79–82 (2005).

32. Pinto, B., Henriques, A. C., Silva, P. M. A. & Bousbaa, H. Three-dimensional spheroids as in vitro preclinical models for cancer research. Pharmaceutics vol. 12 1–38 Preprint at 10.3390/pharmaceutics12121186 (2020).

33. Lee, R. Y. et al. Application of Artificial Intelligence to In Vitro Tumor Modeling and Characterization of the Tumor Microenvironment. Advanced Healthcare Materials vol. 12 Preprint at 10.1002/adhm.202202457 (2023).

34. Linsley, J. W. et al. Superhuman Cell Death Detection with Biomarker-Optimized Neural Networks. Sci. Adv vol. 7 https://www.science.org (2021).

35. Carnevali, F. et al. Advancements in Cancer Research: 3D Models, Single-Cell, and Live-Cell Techniques for Better Insights. Advanced Therapeutics Preprint at 10.1002/adtp.202400351 (2024).

36. Langhans, S. A. Using 3D in vitro cell culture models in anti-cancer drug discovery. Expert Opin Drug Discov 16, 841–850 (2021).

