## Supplementary file for "A Deep Learning Pipeline for Morphological and Viability Assessment of 3D Cancer Cell Spheroids"

### Supplimetry Material-

**Supplementary Fig.1 - MAE per fold (K=10)**

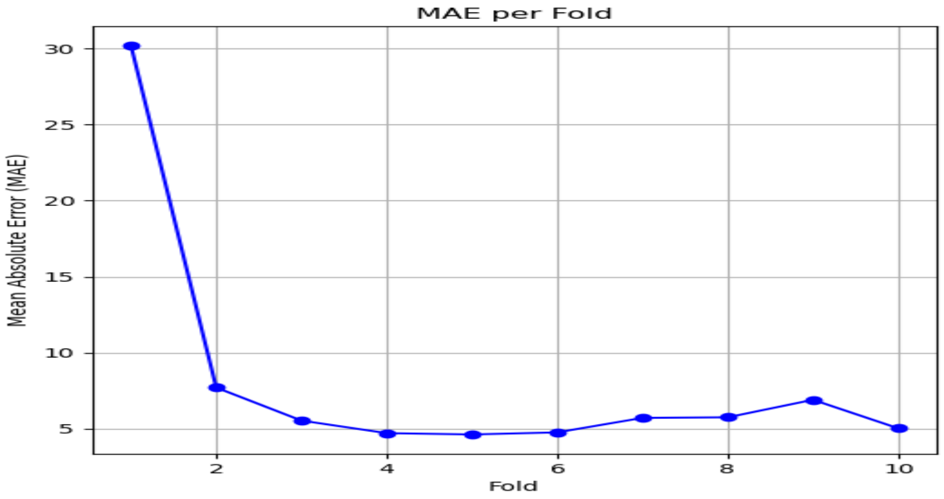


**Supplementary Fig.2- Training and Validation MAE Across Folds (K=10)**

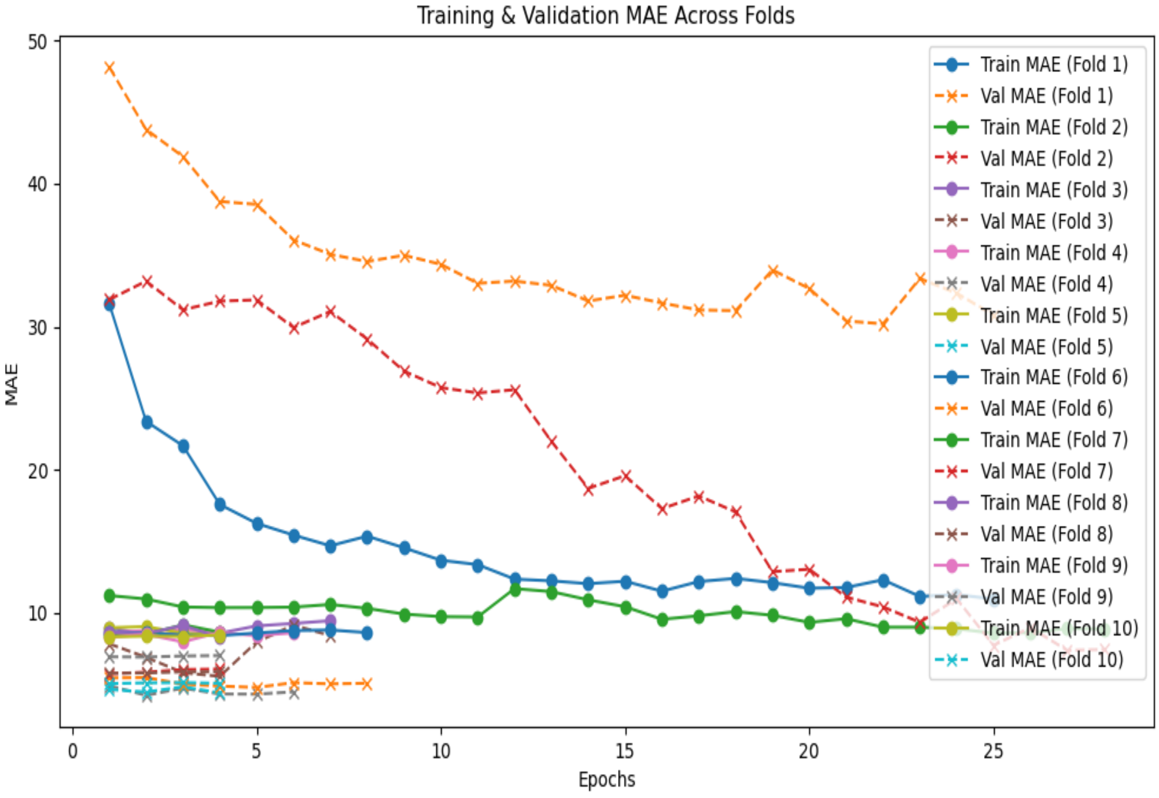

**Supplementary Fig.3- Training and Validation Loss Across Folds (K=10)**

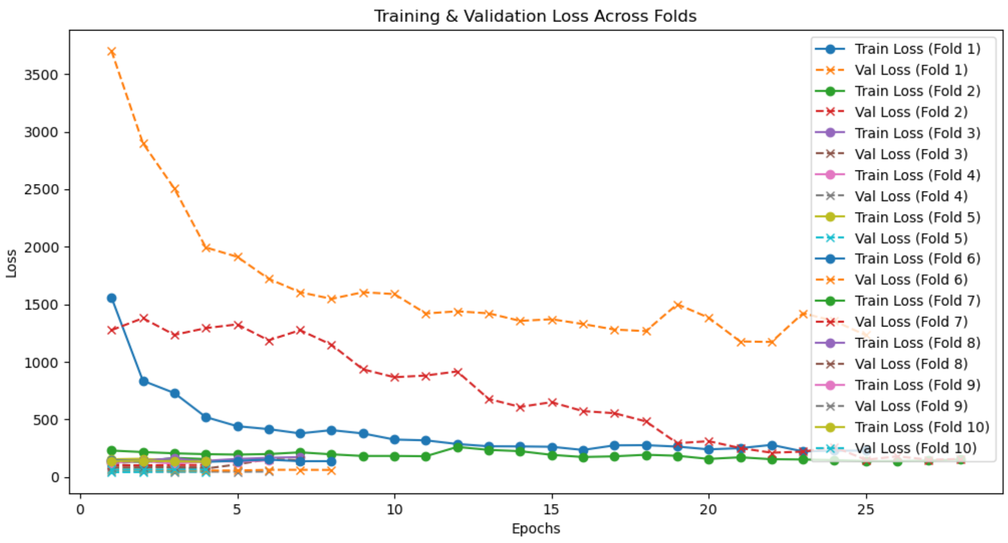


**Supplementary Fig.4 -U-Net Model Accuracy Evaluation Matrix**

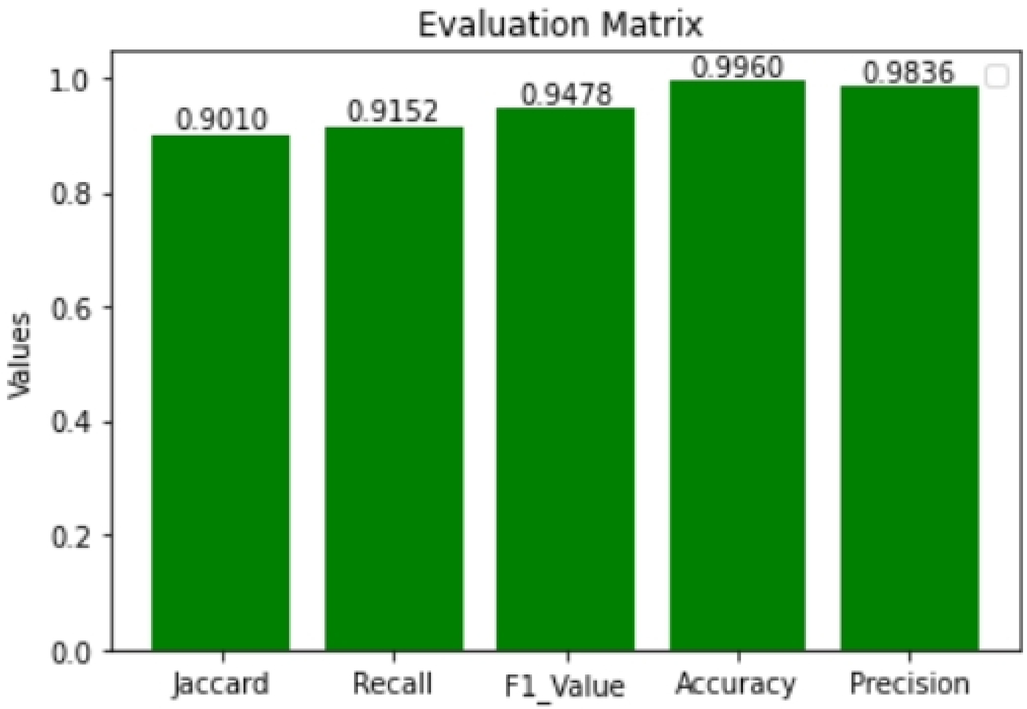
